# Adiposity-Independent Effects of Aging on Insulin Sensitivity and Clearance in Humans and Mice

**DOI:** 10.1101/333997

**Authors:** Nicole Ehrhardt, Jinrui Cui, Sezin Dagdeviren, Suchaorn Saengnipanthkul, Helen S. Goodridge, Jason K. Kim, Louise Lantier, Xiuqing Guo, Yii-Der I. Chen, Leslie J. Raffel, Thomas A. Buchanan, Willa A. Hsueh, Jerome I. Rotter, Mark O. Goodarzi, Miklós Péterfy

## Abstract

**Aims/hypothesis:** Aging is associated with impaired insulin sensitivity and increased prevalence of type 2 diabetes. However, it remains unclear whether aging-related insulin resistance is due to age *per se*, or increased adiposity associated with advanced age. In the present study, we investigate the impact of aging on insulin sensitivity independent of changes in body composition.

**Methods:** Cohorts of C57BL/6J male mice at 4-8 months of age (‘young’) and 18-27 mo (‘aged’) exhibiting similar body composition were characterized with static (plasma glucose and insulin levels) and dynamic (glucose and insulin tolerance tests) measures of glucose metabolism on chow and high-fat diets. Insulin sensitivity was assessed by hyperinsulinemic-euglycemic clamp analysis. The relationship between aging and insulin resistance in humans was investigated in 1,250 non-diabetic Mexican-American individuals who underwent hyperinsulinemic-euglycemic clamps.

**Results:** In mice with similar body composition, age had no detrimental effect on plasma glucose and insulin levels. However, aged mice demonstrated mildly, but reproducibly, improved glucose tolerance on both chow and high-fat diets due to increased glucose-stimulated insulin secretion. Moreover, hyperinsulinemic-euglycemic clamps revealed impaired insulin sensitivity and reduced insulin clearance in aged mice on both diets. Consistent with results in the mouse, age remained an independent determinant of insulin resistance after adjustment for body composition in Mexican-Americn males. Advanced age was also associated with diminished insulin clearance, but this effect was dependent on increased BMI.

**Conclusions/interpretation:** This study demonstrates for the first time that aging *per se* impairs insulin sensitivity independent of adiposity in mice and humans. These results raise the possibility that the pathogenetic mechanisms of age-related and obesity-associated insulin resistance are distinct.

**Abbreviations:**

BAI: body adiposity index
GEE: generalized estimating equations HF high-fat
IQR: interquartile range
MCRI: metabolic clearance rate of insulin
T2D: type 2 diabetes

## INTRODUCTION

The negative influence of aging on insulin sensitivity and the prevalence of type 2 diabetes (T2D) has long been recognized [1]. However, the pathogenetic mechanisms responsible for the age-related deterioration of glucose metabolism remain incompletely understood [2, 3]. A principal difficulty is that aging is associated with numerous adverse physiological changes (e.g. altered body composition, reduced physical activity, sarcopenia, inflammation), which makes it difficult to decipher the role of aging itself from the impact of age-related comorbidities. In particular, as adiposity is a chief determinant of insulin sensitivity, the age-related accumulation of fat mass is a major confounder in studies investigating the metabolic effects of aging [4]. For example, whereas aging-associated declines in glucose tolerance and insulin sensitivity are well documented in the mouse, previous reports are based on comparisons between young and old mice with different body composition [5-8]. Thus, the impact of age *per se* on insulin sensitivity in the mouse is currently unknown.

A substantial body of evidence indicates that insulin sensitivity also declines with advanced age in humans [9-16]. To discriminate between the effects of age versus age-associated changes in body composition, previous studies employed young and elderly cohorts matched for adiposity, or adjusted for this trait in population-based cohorts. Investigations in predominantly non-Hispanic white cohorts consistently failed to demonstrate an effect of aging on insulin sensitivity after adjustment for altered body composition suggesting that increased adiposity, and not age *per se*, is responsible for age-associated deterioration of glucose tolerance in this population [15-20]. In contrast, in a Japanese population-based cohort, age remained positively correlated with insulin resistance even after adjusting for fat content [21]. Remarkably, adjustment for adiposity revealed age-related improvement in insulin sensitivity in other cohorts with predominantly African-American participants [22-24]. Taken together, these observations suggest that genetic and/or other ethnicity-specific determinants may play a role in age-related changes in insulin action and raise the possibility that impaired glucose homeostasis associated with aging may have different etiologies in different populations.

In the present study, we investigate whether aging-related insulin resistance is a direct effect of age, or a consequence of increased adiposity associated with aging in mouse and human subjects. To explore the adiposity-independent effects of aging on insulin action, we assess insulin sensitivity in young and aged mice with similar body composition for the first time. In human studies, we address this question in Mexican-Americans, a population in which the metabolic impact of aging has not been invetigated before despite their strong predisposition to diabetes [25].

## METHODS

### Mice

Young and aged male C57BL/6J mice were purchased from The Jackson Laboratory (Bar Harbor, ME) at 2-3 and 12 months of age, respectively, and maintained at Cedars-Sinai Medical Center until analysis at ages indicated in the figure legends. We used several approaches to ensure that only apparently healthy aged mice were included in our study. First, body weights were measured biweekly and mice showing weight loss at two successive time points were excluded from phenotyping. Second, complete blood cell count was evaluated with a HEMAVET 950 FS counter (Drew Scientific, FL) to identify and exclude mice with lymphomas, a prevalent neoplasm in aging C57BL/6 mice [26]. Finally, after phenotypic characterization, aged mice underwent necropsy and mice with visible tumors and enlarged organs were excluded from analysis. Animals were maintained under specific pathogen-free (SPF) conditions on a 14-h light/10-h dark cycle and fed *ad libitum* Laboratory Rodent Diet 5053 (LabDiet) or a high-fat (HF) diet (D12492; 60 kcal% from lard; Research Diets) for 9 weeks with free access to water. Experimental procedures were approved by the Institutional Animal Care and Use Committees at Cedars-Sinai Medical Center, Western University, University of Massachusetts and Vanderbilt University.

### Mouse phenotyping

Body composition was determined by quantitative magnetic resonance analysis (EchoMRI, TX). Blood glucose concentrations were measured with a OneTouch portable glucose meter (LifeScan, CA). Plasma insulin and C-peptide were assayed by ELISA (ALPCO, NH). In glucose tolerance tests (GTT), overnight fasted mice were retro-orbitally injected with 1.5 g/kg (chow diet) or 1 g/kg (HF diet) glucose followed by blood glucose measurements. Insulin tolerance tests (ITT) were performed in *ad libitum* fed mice by retro-orbital injection of 0.8 U/kg (chow diet) or 1.2 U/kg (HF diet) insulin followed by blood glucose measurements. Hyperinsulinemic-euglycemic clamping of mice was performed at the National Mouse Metabolic Phenotyping Centers (MMPC) at the University of Massachusetts Medical School and Vanderbilt University (Supplemental Material). The metabolic clearance rate of insulin (MCRI) was calculated as insulin infusion rate divided by the mean plasma insulin concentration at steady state [27]. The calculation of MCRI assumes that endogenous insulin secretion is completely suppressed in the setting of hyperinsulinemic clamp, as has been demonstrated in several species [28-30].

### Human study subjects and phenotyping

The current study was conducted in 1,250 non-diabetic subjects from two independent family-based cohorts recruited from the Los Angeles area, the Hypertension-Insulin Resistance (HTN-IR) and the Mexican-American Coronary Artery Disease (MACAD) cohorts. HTN-IR consists of Los Angeles Hispanic-American families ascertained via a proband with essential hypertension [31]. MACAD participants were drawn from adult offspring of probands with coronary artery disease and their spouses [32]. Participants were free of major medical illness and none were taking glucocorticoids or antihyperglycemic agents that could affect glucose homeostasis when they were phenotyped. All subjects gave written informed consent prior to participation.

### Human phenotyping

We studied 579 subjects from HTN-IR and 671 MACAD participants who had undergone phenotyping with the hyperinsulinemic-euglycemic clamp as described previously [33]. The glucose infusion rate (M value) during the last 30 minutes of steady-state glucose and insulin levels during the clamp reflects glucose uptake by all tissues of the body (mainly insulin-mediated glucose uptake in muscle) and is directly correlated with tissue insulin sensitivity [34]. The insulin sensitivity index (M/I) was calculated as M divided by the steady state plasma insulin level (I). The MCRI was calculated as the insulin infusion rate divided by the insulin concentration during the steady state of the euglycemic clamp, as previously described [34, 35].

Body adiposity index (BAI) was calculated as (hip circumference in centimeters)/(height in meters)^1^.^5^-18) [36]. BMI was calculated as weight in kilograms divided by height in meters squared.

All studies were approved by Human Subjects Protection Institutional Review Boards at UCLA, the University of Southern California and Cedars-Sinai Medical Center.

### Statistical analysis

In the mouse data, normally distributed values are expressed as mean ± SEM and non-normally distributed values are represented by the median and interquartile range (IQR). Comparisons between two groups were performed by Student’s t-test or Mann-Whitney U test. GTT and ITT results were analyzed by 2-way repeated measures ANOVA and Holm-Sidak *post hoc* tests. ANOVA and correlation analyses of non-normally distributed variables were performed after transformation or on ranks. Outlier data points were defined as those falling outside 2xIQR. Based on this criterion, 2 extreme values were excluded from analysis in the entire study (1 young and 1 aged mouse in Figure 2G). Differences at p<0.05 were considered statistically significant.

**Figure 1:**
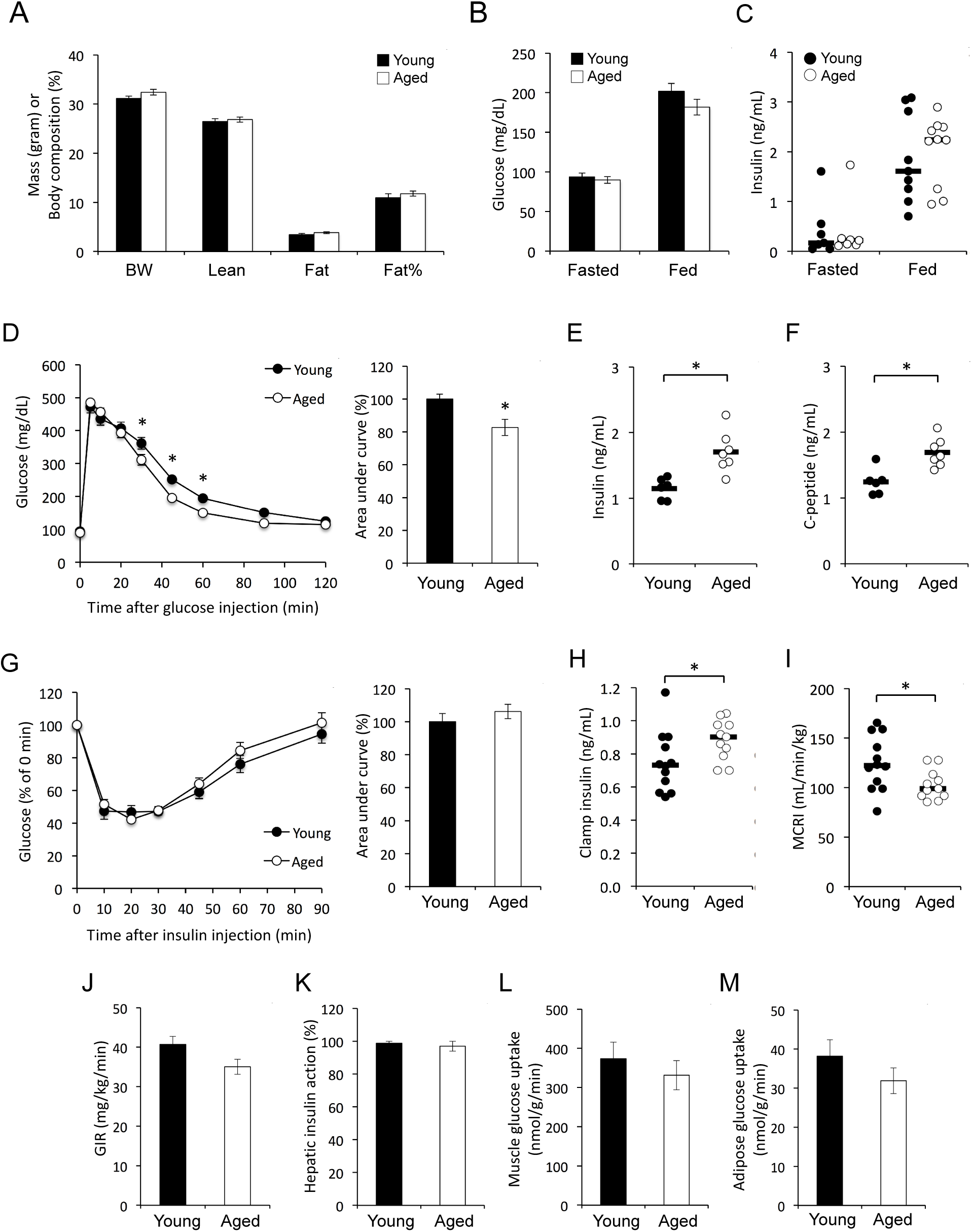
Aging is associated with improved glucose tolerance, decreased insulin clearance and impaired insulin sensitivity in male C57BL/6J mice on a chow diet. *A*: Body weight (BW) and composition of young (4-5 months) and aged (18-21 months) mice (n=9 mice per group). Vertical axis shows mass in grams for BW, Lean and Fat, or body composition as percentage of fat for Fat%. *B*: Blood glucose levels in overnight fasted and *ad libitum* fed mice (n = 9 mice per group). *C*: Plasma insulin levels in overnight fasted and *ad libitum* fed mice (horizontal bars represent median, n = 7-10 mice per group). *D*: Glucose tolerance test (n = 9 mice per group). *P < 0.05, comparison by 2-way repeated measures ANOVA (left panel) or Student’s *t* test (right panel). *E*: Plasma insulin levels 15 minutes after glucose injection (horizontal bars represent mean, n = 6-7 mice per group). *P < 0.05, comparison by Student’s *t*-test. *F*: Plasma C-peptide levels 15 minutes after glucose injection (horizontal bars indicate mean, n = 6-7 mice per group). *P < 0.05, comparison by Student’s *t*-test. *G*: Insulin tolerance test (n=9 mice per group). *H*: Plasma insulin levels during hyperinsulinemic-euglycemic clamp in young (8 months) and aged (27 months) mice (horizontal bars represent median, n = 11-12 mice per group). *P < 0.05, comparison by Mann-Whitney U test. *I*: Metabolic clearance rate of insulin (MCRI) during the clamp (horizontal bars represent median, n = 11-12 mice per group). *P < 0.05, comparison by Mann-Whitney U test. *J*: Glucose infusion rate (GIR) during the clamp (n = 11-12 mice per group). *K*: Suppression of hepatic glucose production during the clamp (n = 11-12 mice per group). Glucose uptake in skeletal muscle (*L*) and adipose tissue (*M*) during the clamp (n = 11-12 mice per group). Data are presented as mean ± SEM unless stated otherwise.

**Figure 2:**
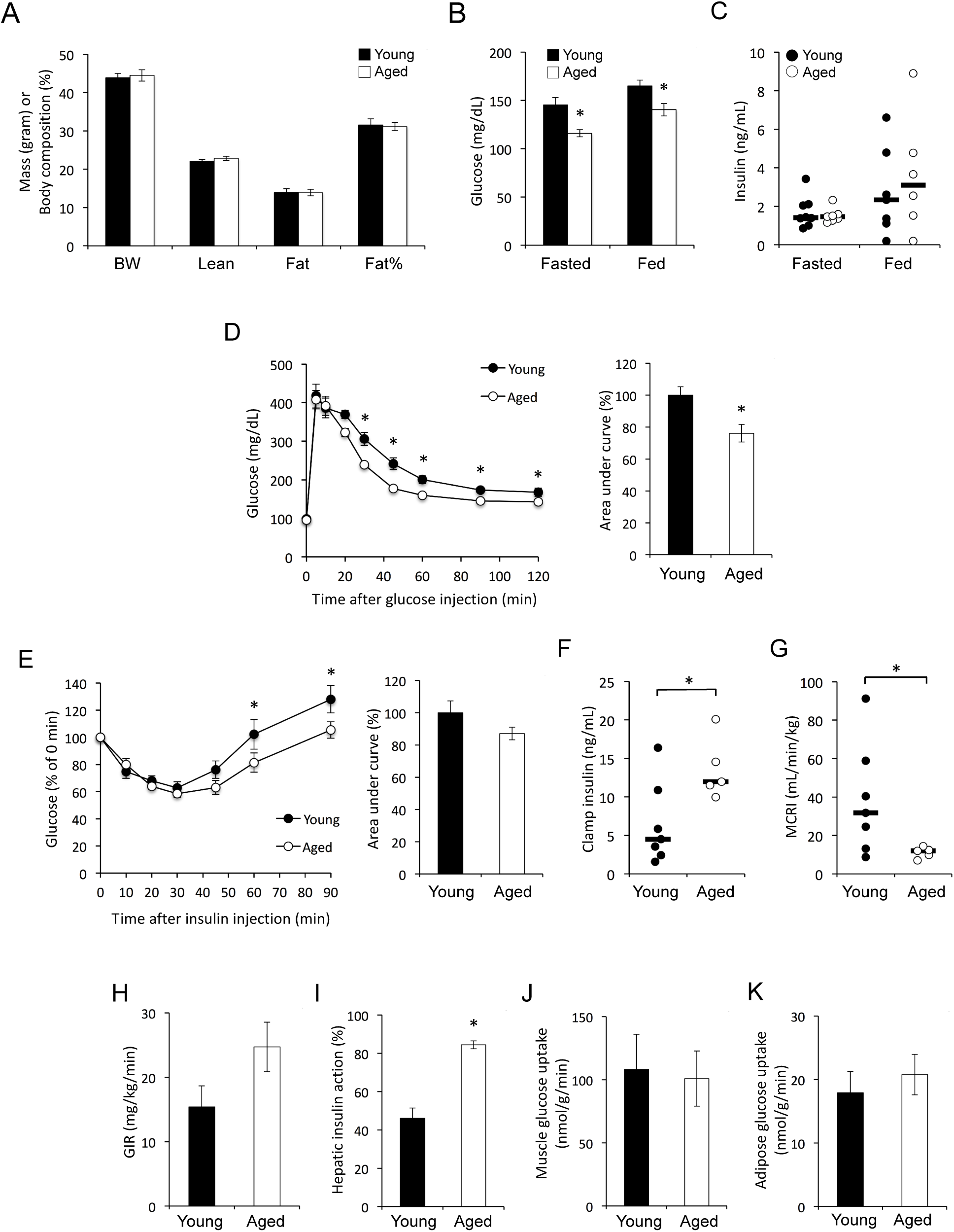
Aging is associated with improved glucose tolerance, decreased insulin clearance and impaired insulin sensitivity in male diet-induced obese C57BL/6J mice. *A*: Body weight (BW) and composition of young (6 months) and aged (24 months) mice (n = 8 mice per group) after 9 weeks of HF diet feeding. Vertical axis shows mass in grams for BW, Lean and Fat, or body composition as percentage of fat for Fat%. *B*: Blood glucose levels in 5-hour fasted and *ad libitum* fed mice (n = 6-8 mice per group). *P < 0.05, comparison by Student’s *t*-test. *C*: Plasma insulin levels in overnight fasted and *ad libitum* fed mice (horizontal bars represent median, n = 5-8 mice per group). *D*: Glucose tolerance test (n = 8 mice per group). *P < 0.05, comparison by 2-way repeated measures ANOVA (left panel) or Student’s *t* test (right panel). *E*: Insulin tolerance test (n = 8 mice per group). *P < 0.05, comparison by 2-way repeated measures ANOVA. *F*: Plasma insulin levels during hyperinsulinemic-euglycemic clamp (horizontal bars represent median, n = 5-7 mice per group). *P < 0.05, comparison by Mann-Whitney U test. *G*: Metabolic clearance rate of insulin (MCRI) during the clamp (horizontal bars represent median, n = 5-7 mice per group). *P < 0.05, comparison by Mann-Whitney U test. *H*: Glucose infusion rate (GIR) during the clamp (n = 6-8 mice per group). *I*: Suppression of hepatic glucose production during the clamp (n = 5-7 mice per group). *P < 0.05, comparison by Student’s *t*-test. Glucose uptake in skeletal muscle (*J*) and adipose tissue (*K*) during the clamp (n = 6-8 mice per group). Data are presented as mean ± SEM unless stated otherwise.

In the human data, log-transformed (BMI, fasting glucose, fasting insulin) or square root-transformed (M value, M/I, MCRI) trait values were used to normalize the distribution for statistical analyses. Generalized estimating equations (GEE) were used to assess the relationships between pairs of traits (univariate analyses) or joint effects of multiple traits (multivariate analyses) on M, M/I, or MCRI, adjusting for familial relationships. The weighted GEE1 [37] was computed assuming an exchangeable correlation structure and using the sandwich estimator of the variance to account for familial correlation present in family data. GEE was used to derive standardized regression coefficients, which in any one regression equation are measured on the same scale, with a mean of zero and a standard deviation of one. They are then directly comparable to one another, with the largest coefficient indicating which independent variable has the greatest association with the dependent variable.

## RESULTS

### Aging is Associated with Improved Glucose Tolerance, and Reduced Insulin Sensitivity and Clearance in Mice

Several previous studies investigating the metabolic impact of aging documented reduced glucose tolerance and insulin resistance in old mice (14-26 months of age) relative to young (1-3 months) mice [5-8]. However, as body fat content increases during the first few months of life, these studies were confounded with different body compositions in old vs young mice and the metabolic effects of age *per se* remained unclear [5-8]. As body weight and adiposity reach a plateau in mature adult (∽4 months) mice [38, 39], we reasoned that using mouse cohorts beyond this age would allow metabolic characterization without the confounding effects of altered body composition. Indeed, mature adult C57BL/6J mice at 4-5 months of age (from here on referred to as ‘young’) exhibited body weight, lean mass, fat mass and percent fat content that were indistinguishable from 18-21-month-old (‘aged’) mice (Fig. 1A). Furthermore, young and aged mice maintained similar fasting and *ad libitum* fed glucose and insulin levels (Fig. 1B and 1C). Intravenous glucose tolerance tests (GTT) revealed that aged mice were slightly, but significantly more glucose tolerant than young mice (Fig. 1D). This was due, at least in part, to increased glucose-stimulated insulin secretion in aged mice, as both insulin and C-peptide levels were elevated in this group 15 min after glucose injection (Fig. 1E and 1F), consistent with previous studies [40]. In insulin tolerance tests (ITT), young and aged mice demonstrated similar glucose response to intravenously-injected insulin (Fig. 1G). To confirm these results, we characterized a second group of young (6-month-old) and aged (25-month-old) mice with similar body composition (Supplemental Fig. 1A). Consistent with results obtained in the initial cohort, aged mice exhibited improved glucose tolerance (Supplemental Fig. 1B), but unchanged insulin tolerance (Supplemental Fig. 1C) relative to young mice.

We further investigated the metabolic impact of aging by assessing glucose homeostasis in a third cohort of young (8-month-old) and aged (27-month-old) male mice by hyperinsulinemic-euglycemic clamp analyses. Body weight and composition, fasting plasma glucose and insulin levels, and the rate of hepatic glucose production were indistinguishable between young and aged mice in the basal state (Supplemental Table 1). However, despite identical rates of insulin infusion, aged mice exhibited elevated plasma insulin levels (Fig. 1H) and a lower rate of insulin clearance from the circulation (Fig. 1I) during the clamp. No significant differences were observed in glucose infusion rate, whole body glucose turnover, suppression of hepatic glucose production, and glucose uptake into muscle and adipose tissue under clamp conditions (Fig. 1J-M, Supplemental Table 1, Supplemental Fig. 2A-B). Based on the observation of similar insulin action despite higher insulin levels in aged animals, we conclude that aging is associated with increased hepatic, muscle and adipose insulin resistance in male mice independent of changes in adiposity.

**Table 1.** Clinical characteristics and their correlation with age in Mexican–Americans. The cohort consists of 42.8% males and 57.2% females. IQR, interquartile range; BMI, body mass index; BAI, body adiposity index; M, glucose disposal rate in the final 30 min of euglycemic clamp normalized by body surface area; M/I, M value divided by steady-state insulin concentration during the clamp; MCRI, metabolic clearance rate of insulin. p<0.05 values appear in bold.

|  | n | Median | IQR | r | p |
| --- | --- | --- | --- | --- | --- |
| Age [years] | 1250 | 34 | 15 |  |  |
| BMI [kg/m<sup>2</sup>] | 1250 | 28.2 | 6.5 | 0.188 | **<0.0001** |
| BAI [%] | 1250 | 31.8 | 8.7 | 0.199 | **<0.0001** |
| Fasting plasma glucose [mg/dl] | 1246 | 92.0 | 12.0 | 0.213 | **<0.0001** |
| Fasting plasma insulin [μU/ml] | 1194 | 12.0 | 8.2 | 0.017 | 0.568 |
| M [mg/min/m<sup>2</sup>] | 1250 | 223 | 150 | -0.205 | **<0.0001** |
| M/I [mg x ml/min/m<sup>2</sup>/mU] | 1191 | 1.72 | 1.43 | -0.192 | **<0.0001** |
| MCRI [ml/m<sup>2</sup>/min] | 1191 | 459 | 135 | -0.062 | **0.020** |

### Aging is Associated with Improved Glucose Tolerance, but Impaired Muscle and Adipose Insulin Sensitivity and Reduced Insulin Clearance in Diet-Induced Obese Mice

The results in lean mice prompted us to investigate whether diet-induced metabolic stress may exacerbate aging-related insulin resistance. To test this hypothesis, we assessed the impact of aging on glucose homeostasis in the context of diet-induced obesity by characterizing young (6-month-old) and aged (24-month-old) male C57BL/6J mice fed a HF diet (60 kcal% from fat) for 9 weeks. At the time of metabolic analysis young and aged mice had similar body weights and composition (Fig. 2A). Unexpectedly, aged mice maintained lower fasting and fed glucose levels (Fig. 2B) despite similar plasma insulin concentrations (Fig. 2C). Consistent with results obtained in lean mice, obese aged mice also demonstrated improved glucose tolerance relative to young animals (Fig. 2D). Furthermore, assessment of insulin tolerance by ITT revealed that exogenous insulin had a greater glucose-suppressing effect in aged mice at later time-points of the assay, although the difference in areas under the curve did not reach statistical significance (Fig. 2E). To evaluate insulin sensitivity directly, we performed hyperinsulinemic-euglycemic clamp experiments. Consistent with results on chow diet, obese aged mice maintained significantly higher insulin levels (Fig. 2F) and demonstrated reduced insulin clearance (Fig. 2G) during the clamp. Despite elevated plasma insulin concentration in aged mice, glucose infusion rate, whole body glucose turnover and glucose uptake in muscle and adipose were similar in young and aged mice (Fig. 2H-K, Supplemental Table 2, Supplemental Fig. 2C-D). However, insulin-mediated suppression of hepatic glucose production was significantly greater in aged mice (Fig. 2I). Taken together, our results indicate that in the metabolic context of diet-induced obesity, aging is associated with improved glucose tolerance, reduced insulin clearance, and impaired muscle and adipose insulin sensitivity in male mice.

**Table 2.** Multivariate correlations between insulin sensitivity and age in Mexican-Americans. Mand **M/1** are dependent variables; Age and measures of body composition are independent variables. Standardized coefficients() and p values for Age are shown. Mand **M/1,** and **BMI** were square root or log-transformed, respectively, before analysis. n, number of subjects. p<0.05 values appear in bold.

| Independent variables | Males |  |  |  |  |  | Females |  |  |  |  |  |
| --- | --- | --- | --- | --- | --- | --- | --- | --- | --- | --- | --- | --- |
|  | M |  |  | M/I |  |  | M |  |  | M/I |  |  |
| | n | $\beta$ | p | n | $\beta$ | p | n | $\beta$ | p | n | $\beta$ | p |
| Age | 535 | -0.279 | **<0.0001** | 505 | -0.260 | **<0.0001** | 715 | -0.140 | **<0.0001** | 686 | -0.133 | **<0.001** |
| Age, BMI | 535 | -0.205 | **<0.0001** | 505 | -0.186 | **<0.0001** | 715 | -0.065 | **0.044** | 686 | -0.038 | 0.241 |
| Age, BAI | 535 | -0.221 | **<0.0001** | 505 | -0.197 | **<0.0001** | 715 | -0.054 | 0.116 | 686 | -0.031 | 0.373 |

### Sex-specific Effect of Aging on Insulin Sensitivity and Clearance in Mexican-Americans

To explore the relationship between age, insulin resistance, insulin clearance and adiposity in humans, we performed correlation analyses in a Mexican-American cohort of 1,250 participants free of diabetes and overt metabolic disease. The age range of the cohort is 7 decades (16-87 years) and insulin sensitivity has been assessed by hyperinsulinemic-euglycemic clamp. Univariate correlation analyses revealed aging-associated changes in glucose homeostasis including elevated fasting plasma glucose levels, reduced insulin sensitivity (M and M/I) and decreased MCRI (Table 1). To assess body composition, we used BMI and BAI, an alternative surrogate measure of fat content more closely tracking percent fat content in Mexican-Americans [36]. Both BMI and BAI exhibited strong positive correlations with age indicating fat accumulation during aging in this population (Table 1).

Sex-specific analysis demonstrated significant negative correlations between age and insulin action in both males and females with notably weaker effects in the latter (Table 2). To separate the contributions of age and adiposity on insulin action, we performed multivariate correlation analyses with measures of insulin sensitivity as dependent variables, and age and body composition as independent variables. In males, age remained strongly correlated with insulin resistance even after adjustment for changes in body composition (Table 2). In contrast, controlling for adiposity reduced or abolished the correlation between age and insulin action in females, depending on the measure of insulin sensitivity used (Table 2). While the effect of age was relatively modest in the multivariate models tested (Table 2), BMI and BAI had robust effect sizes and remained strongly negatively correlated with insulin sensitivity after controlling for age in both sexes (Supplemental Table 3). Taken together, these results demonstrate an age-dependent decline of insulin sensitivity in Mexican-American males and females. In females, this is principally due to aging-associated increases in fat mass, whereas age is an independent determinant of insulin resistance in males.

**Table 3.** Multivariate correlations between insulin clearance and age in Mexican-Americans. MCRI is the dependent variable; Age and measures of body composition are independent variables. Standardized coefficients (β) and p values for Age are shown. MCRI and BMI were square root or log-transformed, respectively, before analysis. n, number of subjects. p<0.05 values appear in bold.

| Independent variables | Males |  |  | Females |  |  |
| --- | --- | --- | --- | --- | --- | --- |
| | n | $\beta$ | p | n | $\beta$ | p |
| Age | 505 | -0.104 | **0.008** | 686 | -0.033 | 0.335 |
| Age, BMI | 505 | -0.068 | 0.112 | 686 | 0.029 | 0.422 |
| Age, BAI | 505 | -0.069 | 0.107 | 686 | 0.023 | 0.527 |

Sex-specific analysis of insulin clearance revealed that the age-related decline observed in the full cohort (Table 1) was exclusively due to males, as significant correlation between age and MCRI could not be detected in females (Table 3). Furthermore, multivariate analysis indicated that the aging-associated decrease of insulin clearance in males was due to increased body fat content and not age *per se* (Table 3).

## DISCUSSION

The association of aging with impaired insulin resistance has been well established in mice [5-8]. However, as body composition changes with normal aging, previous studies have been unable to disentangle the effects of aging per se from those of increased adiposity. In the present study, we address this problem for the first time by characterizing mice of different age, but similar body composition. We took advantage of the fact that body weight and fat content in C57BL/6 mice reach a plateau at the mature adult stage (∽4 months) and remain unchanged thereafter [38, 39]. Thus, using ‘young’ mouse cohorts at >4 months of age allowed us to investigate the adiposity-independent effects of aging on glucose metabolism for the firat time.

When mice with similar body compositions were compared, advanced age had no detrimental impact on static (i.e. plasma glucose and insulin levels) or dynamic measures (i.e. GTT, ITT) of glucose metabolism. In fact, we found that aged mice exhibited slightly improved glucose tolerance, an observation that was replicated in three independent mouse cohorts on two different diets. Consistent with these results, improved glucose tolerance in aging C57BL/6J mice has also been previously reported [40]. Importantly and similar to the present study, Leiter et al. used ‘young’ mice at the relatively advanced age of 4.5-to 5 month, which likely diminished differences in body composition between young and old mice, although adiposity was not directly assessed in that study [40]. It has also been demonstrated that aging is associated with increased pancreatic islet area and insulin content, but unaffected β-cell sensitivity to glucose [40]. These results are consistent with our observation of elevated C-peptide levels after glucose administration in aged mice and suggest that elevated glucose-stimulated insulin secretion contributes to their improved glucose tolerance. Another likely contributing factor is reduced insulin clearance in old mice. We hypothesize that the interaction between increased production and reduced clearance of insulin may explain the apparent over-compensation for insulin resistance in aged mice. In conclusion, while glucose intolerance is a hallmark of aging [7, 41], our results demonstrate that this is not an inevitable consequence of advanced age, but most likely due to increased adiposity associated with aging in C57BL/6J mice.

We assessed the impact of aging on insulin sensitivity by hyperinsulinemic-euglycemic clamps in young and aged mice with similar body compositions. Despite similar rates of insulin infusion, aged mice exhibited significantly higher plasma insulin concentration during the clamp. This observation held true for both lean mice maintained on a normal chow diet and diet-induced obese mice on a HF-diet. In lean mice, measures of whole-body and tissue insulin action were similar in young and aged animals. Together with elevated insulin levels in the latter, these data demonstrate for the first time that aging *per se* impairs insulin sensitivity in C57BL/6J mice.

We initially hypothesized that the metabolic stress associated with HF diet feeding would exacerbate aging-related impairments in glucose homeostasis. However, our results do not support this hypothesis. In fact, obese aged mice maintained lower basal glucose levels and exhibited improved glucose tolerance relative to their young counterparts. Furthermore, we unexpectedly found that aging impacted tissues differently in diet-induced obese mice. Consistent with observations in lean mice, despite elevated clamp insulin levels, insulin action was similar in muscle and adipose tissues of young and aged animals indicating relative insulin resistance in the latter. In contrast, suppression of hepatic glucose production was substantially stronger in obese aged versus young mice. Although the concurrent elevation of insulin levels complicates the interpretation of this observation, our results suggest that hepatic insulin sensitivity may not be adversely impacted by age in the context of diet-induced obesity. Consistent with this conclusion, aging has also been associated with improved hepatic insulin action in humans in a handful of studies where hepatic glucose production was determined, [22, 42].

Most previous human studies focusing predominantly on non-Hispanic white populations led to the consensus view that aging-associated insulin resistance is due to age-related increase in adiposity rather than age *per se* [15-20, 22]. In the current study, we addressed this issue in a cohort of non-Hispanic white subjects for the first time. Consistent with previous reports, insulin sensitivity declined with age in both sexes. However, the relative contributions of adiposity and age to aging-associated insulin resistance were different in the sexes. In Mexican-American females, diminished insulin action was due to increases in adiposity and not age itself, in line with results in Caucasians. In contrast, while elevated adiposity was a major determinant of declining insulin action in males, age also contributed to insulin resistance independently of changes in body composition. A similar conclusion was reached in a recent study in Japanese subjects [21]. Taken together, these results highlight the previously underappreciated role of sex and ethnicity in aging-associated insulin resistance in humans.

Previous studies investigating the impact of aging on insulin clearance yielded mixed results [9, 15, 43-45]. Here we report consistent age-related declines in the rate of insulin removal from the circulation in mouse and human subjects. Nonetheless, the underlying mechanisms appear to be different in the two species. Whereas increased adiposity is the main driver of reduced insulin turnover in aging Mexican-Americans, diminished insulin clearance in aged C57BL/6J mice is independent of body composition. This observation is consistent with the recent demonstration of age-related loss of hepatic endothelial fenestrations and diminished insulin uptake associated with reduced insulin clearance in mice and rats [46]. Our results also raise the question of whether age-related changes in insulin clearance are the consequence of aging *per se* or secondary to insulin resistance. As aging is associated with impaired insulin sensitivity in both Mexican-Americans and the mouse model we used, further studies will be required to discriminate between these possibilities.

In summary, the present study provides evidence in mouse and human populations that aging impairs insulin sensitivity independently of altered body composition. Based on the analysis of a Mexican-American cohort, this conclusion contrasts with previous results obtained in non-Hispanic white populations and suggests a role for genetic factors in age-related metabolic dysfunction. The present work also highlights the impact of sex on insulin resistance, as age was an independent contributor to this trait only in Mexican-American men, but not women. As our mouse study was limited to the characterization of C57BL/6J males, future studies involving females and additional mouse strains will be needed to investigate the role of sex and genetic determinants in age-related changes in insulin sensitivity. An intriguing possibility raised by the present study is that the mechanisms of obesity-and aging-associated insulin resistance may be distinct, a notion consistent with a previous report [47]. This possibility may also have important therapeutic implications for the treatment of T2D in the aging population. As demonstrated in the present study, male C57BL/6J mice represent a suitable animal model to investigate the molecular mechanisms underlying the adiposity-independent effects of aging on insulin resistance.

## Acknowledgements

The authors thank Drs. Maura Rossetti, Xiao Z. Shen, Tuantuan Zhao, Shuang Chen and Sujin Suk for their contributions to this project.

## Funding

This study was supported by National Institutes of Health grants R01-HL088457, R01-DK079888, R01-HL67974, P30-DK063491, P50-HL55005, M01-RR000425, M01-RR000043, UL1-TR000124, U2C-DK093000, U2C-DK059637 and P30-DK020593.

## Duality of Interest

The authors declare no conflict of interest.

## Author Contributions

N.E. conducted experiments, analyzed data and revised the manuscript. J.C. analyzed data. S.D. and S.S. conducted experiments. H.S.G. provided materials and advised experiments. J.K.K. and L.L. supervised experiments, analyzed data and revised the manuscript. X.G., Y.I.C., L.J.R., T.A.B., W.A.H. and J.I.R acquired data. M.O.G. and M.P. conceived of the study design, supervised the study, analyzed data, and revised the manuscript. M.P. wrote the manuscript. All authors approved the final version of the manuscript. M.O.G. and M.P. are guarantors of this work and, as such, had full access to all the data in the study and take responsibility for the integrity of the data and the accuracy of the data analysis.

